# Development of functional corticomuscular connectivity from childhood to adulthood during precision grip with the dominant and non-dominant hand

**DOI:** 10.1101/2020.12.22.423542

**Authors:** Mikkel Malling Beck, Meaghan Elizabeth Spedden, Jesper Lundbye-Jensen

## Abstract

How does the neural control of manual movements mature from childhood to adulthood? Here, we investigated developmental differences in functional corticomuscular connectivity using coherence techniques in 91 individuals recruited from four different age groups covering the age range 8-30y. EEG and EMG were recorded while participants performed a unimanual visual force-tracing task requiring fine control of the force produced in a precision grip with both the dominant and non-dominant hand. Using beamforming methods, we reconstructed source activity from EEG data displaying peak coherence with the active FDI muscle during the task in order to assess functional corticomuscular connectivity. Our results revealed that coherence was greater in adolescents and adults than in children and that the difference in coherence between children and adults was driven by a greater magnitude of descending (cortex-to-muscle) coherence. This was paralleled by the observation of a posterior-to-anterior shift in the cortical sources displaying corticomuscular coherence within the contralateral hemisphere from late adolescence. Finally, we observed that corticomuscular coherence was higher on the non-dominant compared to the dominant hand across age groups. These findings provide a detailed characterization of differences in task-related corticomuscular connectivity for individuals at different stages of typical ontogenetic development that may be related to the maturational refinement of dexterous motor control.

**Key points:**

‐ Fine motor control is gradually refined during human motor development, but little is known about the underlying neurophysiological mechanisms.
‐ Here, we used EEG and EMG to investigate functional corticomuscular connectivity during a precision grip with the dominant and non-dominant hand in 91 typically developed children, adolescents and adults (age range 8-30y).
‐ We show that older adolescents and adults are characterized by greater levels of corticomuscular coherence compared to children and that this is mainly driven by greater magnitudes of descending coherence (cortex-to-muscle). This is paralleled by a more anterior cortical site of coherence in older adolescents and adults compared to younger individuals.
‐ These results help us better understand the ontogenetic development of task-related functional connectivity in sensorimotor networks.

## Introduction

Humans develop their dexterous abilities from childhood though adolescence to adulthood (Forssberg *et al*., 1991, 1992; Dayanidhi *et al*., 2013). Efficient functioning of sensorimotor neural networks is a prerequisite for fine control of the fingers during motor tasks. Development of corticomuscular control mechanisms could potentially contribute to improvement of skilled capacity as we age (Olivier *et al*., 1997), but little is known about the actual neurophysiological mechanisms that subserve improvements in dexterity from childhood to adulthood. e human central nervous system (CNS), including the descending and ascending pathways between the brain and spinal cord, is continuously shaped during ontogenetic development. This is e.g. reflected in gradual increases in white matter integrity in the corticospinal tract (Paus *et al*., 1999; Lebel & Beaulieu, 2011) and an increase in the central conduction velocity of both ascending and descending pathways measured via brain and muscle responses to synchronous activation of a peripheral nerve by electrical stimulation and the corticospinal system using transcranial magnetic stimulation (TMS) of the primary motor cortex (M1), respectively (Müller *et al*., 1991, 1994; Nezu *et al*., 1997). These maturational adaptations can shape the passing and processing of information in functional neural networks (Smith *et al*., 2017) and thereby likely also affect patterns of connectivity between brain and spinal cord. However, little is actually known about developmental differences in task-related functional connectivity in corticomuscular networks. Oscillatory coupling between neuronal populations within the CNS may represent a processing strategy allowing efficient neural interactions (Fries, 2015). During voluntary motor tasks, oscillatory activity in parts of the cerebral cortex dedicated to sensorimotor functions correlates with similar rhythms in the contralateral contracting muscles in adult humans (Conway *et al*., 1995) and non-human primates (Baker *et al*., 1997). These patterns of functional connectivity can be captured by measures of corticomuscular coherence reflecting frequency-domain correlations between brain activity obtained by magneto- or electroencephalography (M/EEG) and muscle activity from electromyographic (EMG) recordings. Coherence between the rhythmic signals of the brain and muscle in the beta band (15-30 Hz) is particularly prominent during steady voluntary muscle contractions (Conway *et al*., 1995; Salenius *et al*., 1997).

Patterns of correlated oscillatory activity in the corticomuscular system has been found to emerge during late childhood and to steadily increase in strength until early adulthood (James *et al*., 2008; Spedden *et al*., 2019*b*). Previous studies have investigated developmental differences in the functional coupling between brain and muscle activity for several different muscles including the hand (Graziadio *et al*., 2010), forearm (James *et al*., 2008) and ankle (Spedden *et al*., 2019*b*) muscles on either right or left side of the body. Levels of coherence have been shown to follow a proximal-distal gradient, with higher magnitudes observed for the latter (Ushiyama *et al*., 2010). Additionally, corticomuscular coherence during steady-state muscle contractions have been associated with greater control over motor output, as evidenced by smaller fluctuations in the force produced (Kristeva *et al*., 2007) and less variable muscle activity (Witte *et al*., 2007). Collectively, these studies demonstrate a gradual strengthening of oscillatory corticomuscular interactions during development that could relate to the ability to control fine motor output. However, it is currently unknown whether differences exist between the dominant and non-dominant limb and whether this may change during development.

Early studies advocated that corticomuscular coherence is mediated by beta-range oscillations descending from cortex via the corticospinal tract to the spinal cord enslaving the activity of motor neurons (Baker *et al*., 2003; Hansen & Nielsen, 2004), but emerging evidence suggests that this interpretation may be somewhat simplified (Baker, 2007). For example, administration of pharmacological agents can alter the power of cortical beta-range oscillations or corticomuscular coherence selectively; a result that is at odds with the notion of an exclusive descending and efferent origin of coherence (i.e. a pure feedforward system) (Baker & Baker, 2003; Riddle *et al*., 2004). Instead, sensory (ascending) inputs to sensorimotor cortices might also affect levels of corticomuscular coherence (Hansen & Nielsen, 2004; Riddle & Baker, 2005; Witham *et al*., 2011), thus forming a sensorimotor loop between the cortex and periphery that may be relevant for binding of motor commands and their sensory consequences. Measures of coherence capture the statistical association between signal pairs in the frequency domain, quantifying the degree of functional connectivity, but not the direction. In contrast, measures of *directed* connectivity allow a dissection of descending (from cortex to muscle) and ascending (from muscle to cortex) contributions to coherence based on time lags between the two signals (Halliday, 2015). From a developmental perspective, such information is highly relevant, as it may reveal fundamental insights into potential shifts in the neural control mechanisms that are used to guide behavior. In a recent study, we demonstrated that the degree of descending coherence (from cortex to muscle) increased at the expense of ascending coherence (from muscle to cortex) as a function of age from childhood through adolescence. These increased levels of descending coherence could reflect increased reliance and/or efficiency of feedforward control guiding motor behavior with a concomitant reduction in the importance of ascending feedback (Spedden *et al*., 2019*b*). Notably, EEG was recorded from a single electrode in this study and focus was on the control of the ankle muscles. Therefore, developmental differences in directed corticomuscular interactions and the localization of coherent brain sources as well as the potential role of these control mechanisms for force control for the dominant and non-dominant hand have yet to be explored.

Here, we investigated functional corticomuscular connectivity during a unimanual force-tracing task involving steady and precise control of the force generated by the intrinsic hand muscles with the aim of characterizing age-related differences from childhood to adulthood and explore effects of hand dominancy in a large sample of typically developed individuals. Based on recent results (Spedden et al. 2019*b*), we hypothesized that magnitudes of corticomuscular coherence would increase with age for both the dominant and non-dominant hand and that the amount of descending corticomuscular coherence (cortex-to-muscle) would be greater for older compared to younger individuals. Based on the hypothesis relating to age-related differences in coherence - and in directed connectivity particularly - we furthermore hypothesized that there would be a posterior to anterior shift in the spatial localization of cortical sources displaying coherent activity with the active muscles with increasing age. To our knowledge, this is the first study to address this question.

## Materials and methods

### 2.1 Participants

Ninety-eight individuals were recruited to participate in the main experiment. Participants received thorough oral and written information and written consent was obtained from participants (>18y) or their parents (<18y) prior to enrollment. The study was approved by the regional ethical committee (H-17019671) and adhered to the principles set out in the declaration of Helsinki II. EEG and motor performance data from the dominant hand has already been presented in a previous paper, in which we assessed age-related differences in cortico-cortical connectivity and motor performance (Beck et al., in review). Here, we present data on corticomuscular coherence and task performance for the same task from both the dominant and non-dominant hand. Participants’ handedness were determined by the Edinburgh handedness inventory (Oldfield, 1971) and levels of physical development were characterized using a sketched version of the Tanner scale (Marshall & Tanner, 1969, 1970).

### 2.2 Experimental procedure

EEG was acquired while participants performed a tonic force-tracing task with both the dominant and the non-dominant hand. The procedure for each hand was similar. Participants sat in a chair in front of a computer monitor. Participants initially performed three maximal voluntary contractions (MVC). Then, a horizontal target line corresponding to 10% MVC was presented in the middle of the screen in front of the participants. Participants were asked to apply force to a load cell (Dacell, AM210, Dacell Co. Ltd., Korea) located between their index finger and thumb in a precision grip and match the force produced to a horizontal target line (Figure 1AB). The force produced was amplified ×100 and high-pass filtered at 10Hz (Dacell, AM210, Dacell Co. Ltd., Korea) before being digitized at 1000Hz (CED1401, Cambridge Electronic Design Ltd, Cambridge, UK) and fed back to the participant as a real-time trace of the applied force (Figure 1B). The task was performed for ∼120 sec on each hand starting with the non-dominant. This specific task was chosen because periods of steady contraction in the precision grip are characterized by corticomuscular coherence in the beta-range (15-30Hz) in adults. Performance in the force-tracing task was quantified as motor precision and motor variability. Motor precision was defined as the root mean squared error (RMSE) from the horizontal target line (i.e. deviations from ‘optimal’ performance. This score was reverse coded (multiplied by −1) to reflect motor precision (i.e. inverse error). Furthermore, we also quantified variability of motor output by computing the coefficient of variation (CV) of the force produced. In a subsequent control experiment, we recruited 15 adults who were naïve to the task. These individuals completed the same experiment as described above, but with the order of tasks reversed (i.e. the task was performed with the dominant hand first). This was done to account for potential order effects (results are presented in the supplementary results).

**Figure 1.**
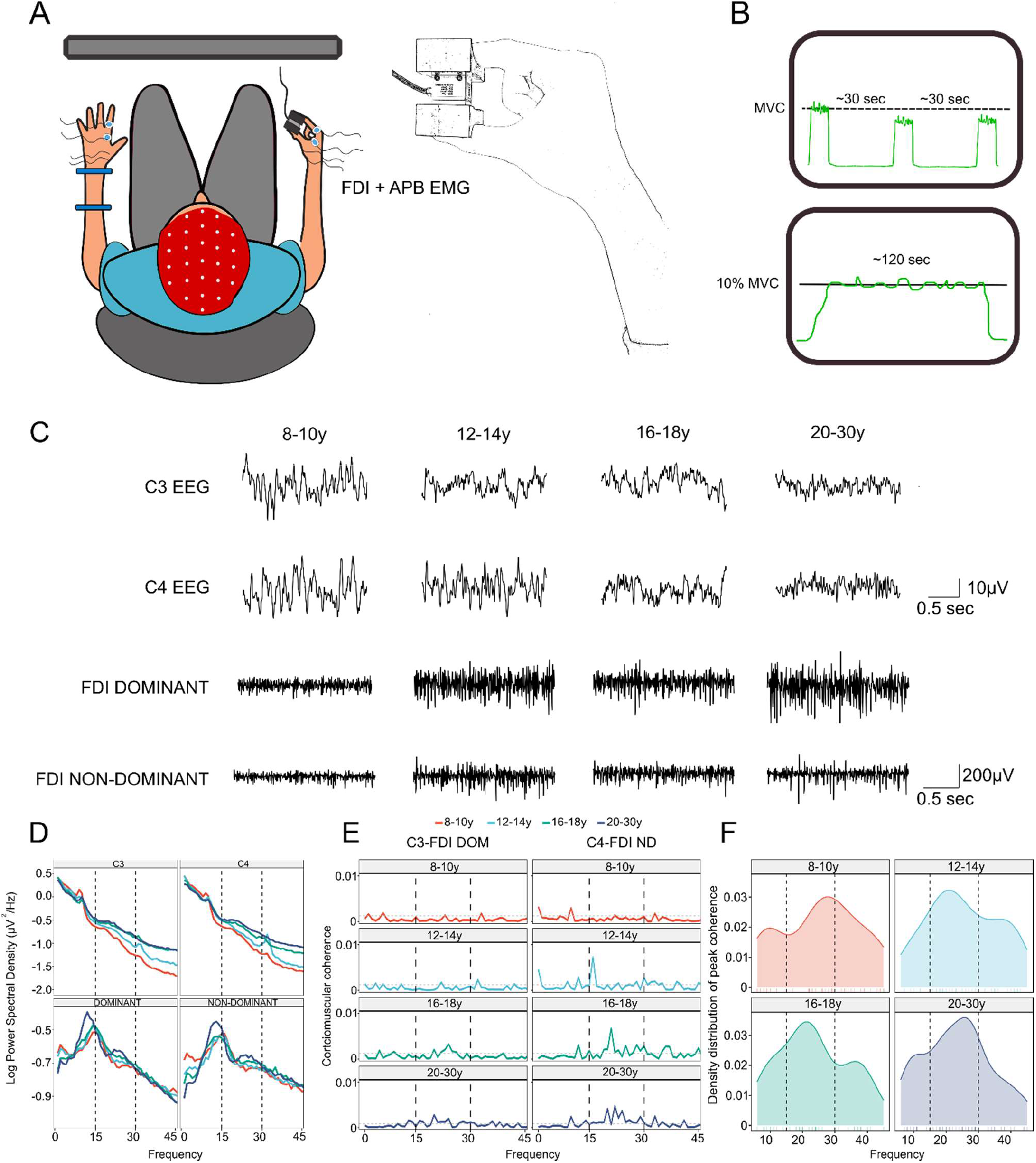
Experimental setup and scalp EEG data. EEG was recorded from 64-channels and EMG was obtained from the FDI and APB muscles of both the dominant and non-dominant hand. Participants initially performed three maximal voluntary contractions (MVC) on the non-dominant hand, before performing the force-tracing task in which participants were asked to maintain a steady force between the index and thumb corresponding to 10% of MVC. Visual feedback was provided as a real-time trace of the applied force (A and B). In C, sensor EEG data and FDI EMG are presented from representative individuals from electrode C3 and C4 (located above primary motor cortex) and the dominant and non-dominant FDI muscles. D presents group-averaged sensor-level power spectral densities from the C3 and C4 electrode alongside the dominant and non-dominant FDI, and in E pooled coherence spectra from each of the four age groups are presented. In F, we present group density plots of the distribution of peak frequency in the 5-45Hz frequency range. The dashed vertical black lines in D, E and F mark the beta-band (15-30Hz). Please note the common legend used for the plots in DEF. Abbreviations: DOM = dominant hand; ND = non-dominant hand; FDI = First dorsal interosseous; APB = Abductor pollicis brevis.

### 2.3 Data acquisition

EEG and EMG were sampled using a BioSemi amplifier system (BioSemi, Amsterdam, The Netherlands) using ActiView software (v7.07) installed on a PC. EEG was obtained from 64 sensors placed in a standard electrode cap with the 10/20 layout (BioSemi, Amsterdam, The Netherlands), and EMG data was obtained from the first dorsal interosseous (FDI) and abductor pollicis brevis (APB) muscle on both the dominant and non-dominant hand (Figure 1A; left). Data was sampled as raw signals at 2048 Hz during the force-tracing task described above. Before performing the task, participants were instructed to keep their neck and face relaxed to avoid excessive muscle activity in the EEG. Furthermore, we instructed participants on keeping the hand not performing the task still and resting on the table. Electrode offsets were <30mV for all EEG sensors prior to initiating the recordings. The common mode sensor (CMS) and the driven right leg (DRL) electrodes provided online referencing.

### 2.4 EEG and EMG preprocessing

EEG and EMG data was preprocessed in EEGLAB (v14.1.1b)(Delorme & Makeig, 2004) in Matlab R2017b. All subsequent steps were performed on files from both the dominant and the non-dominant hand. Raw data files were imported using the Biosig toolbox. Data was visually inspected and time periods in the EEG exhibiting high amplitude fluctuations due to muscle activity were removed. EEG data was separated from EMG data. Then, EEG data was band-pass filtered from 0.5-48Hz and subsequently downsampled to 256 Hz. EEG data was visually inspected and bad channels displaying excessive noise were removed prior to re-referencing sensor activity to average reference. Finally, independent component analysis (ICA) was performed (‘*runica*’ algorithm) to decompose the EEG data into orthogonal components. Components reflecting eye-blinks and/or horizontal eye movements were removed from the data before the removed channels were interpolated using the default spherical interpolation procedure. To limit the total number of analyses, we chose to focus on the EMG data from the FDI and not the APB muscle. This was filtered between 5 and 120 Hz, downsampled to 256 Hz and full-wave rectified. Next, data files were converted to Statistical Parametric Mapping formatted files and epoched in non-overlapping segments using the ‘arbitrary epochs’ option with an epoch-length of 1-sec using SPM12 (v. 7490). Next, default electrode positions were registered. Electrode positions were flipped across the midline in the sagittal plane for left-hand dominant individuals (n=8) to enable group-level comparisons.

### 2.5 Data analysis

Subsequent steps were performed on files from both the dominant and the non-dominant hand: The first descriptive analyses were performed at the sensor level. As a first step, we computed power spectral densities from the C3 and C4 electrode and the FDI EMG from both the dominant and non-dominant hand. Power spectra were computed using a Finite Fourier Transformation (FFT) to represent the signal in the frequency domain. The choice of electrodes was motivated by the fact that corticomuscular coherence generally is topographically localized contralateral to the contracting muscles at electrodes overlying primary motor (M1) regions (Mima & Hallett, 1999).

Next, we aimed to characterize the frequency distribution of corticomuscular coherence. This was done by two complementary approaches. First, we computed coherence spectra for each individual. Coherence between two signals (x,y) at a given frequency C_xy_(f) is represented as the squared magnitude of the cross-spectrum between the two signals (G_xy_) divided by the product of the two auto spectra (G_xx_(f) and G_yy_(f)), with values bound between 0 and 1. Coherence values of zero at a given frequency indicate a total lack of statistical association between the two signals, whereas values of one represent an ideal linear correlation in the frequency-domain. Time series from the C3 and C4 electrodes and the EMG from the FDI muscle were extracted and subjected to this analysis. Power and coherence spectra were visually inspected to assess data quality. Next, individual coherence spectra were pooled on the group-level to provide a summary of group data (Amjad *et al*., 1997). These analyses were performed using the Neurospec tools (v 2.11) (Halliday *et al*., 1995). Second, we also extracted the frequency value where coherence was maximal from the individual spectra and plotted the distribution of peak frequencies for each age group. In accordance with earlier studies, these descriptive analyses revealed that, for all age groups, peak coherence was most commonly observed within the beta band (15-30Hz) (Figure 1EF).

After having established the frequency distribution of corticomuscular coherence at the sensor-level we next turned to localize brain sources that were coherent with muscle activity in the active FDI muscle. We focused our analysis on the beta band because (1) our sensor-level data suggested that this was the most common frequency band in which peak coherence occurred; and (2) previous studies have shown that corticomuscular coherence is particularly pronounced in the beta-range during steady-state motor output (as reviewed in Mima and Hallett 1999; Grosse et al. 2002; van Wijk et al. 2012). Cortical sources that were coherent with muscle activity in the active FDI muscle were identified using Dynamic Imaging of Coherent Sources (DICS) beamforming (Gross *et al*., 2001) implemented in the Data Analysis in Source Space (DAiSS) toolbox in SPM12. DICS uses the cross-spectral density (CSD) matrix combined with a forward model to localize coherent sources of activity in the brain with a peripheral signal (here, the FDI-EMG channel). Beamforming techniques utilize a spatial filter at each point of a search grid of the brain to maximize the source activity and attenuate the activity originating from other locations. These filters are a linear transformation of the lead fields and the CSDs. Lead fields were computed using the Boundary Element Model (BEM) based on the template MRI provided in SPM12 (Litvak *et al*., 2011). Generally, beamforming approaches do not contain prior assumptions about the number and orientation of active (coherent) sources, but instead computes coherence for a ‘full’ source grid comprising the entire brain volume. The source grid resolution was set to 5 mm, and the results were stored as individual SPM volumetric images.

Then, individual images were smoothed using a Gaussian kernel (10×10×10mm) and both group and grand averages were computed from the participants’ individual images. For descriptive comparisons of group-level locations displaying peak coherence we utilized the SPM anatomy toolbox (Eickhoff *et al*., 2005). Using Linear Constrained Minimum Variance (LCMV) beamforming, individual time-series of source activity were extracted as a ‘virtual electrode’ from the grand-average location displaying beta-range peak coherence (MNI coordinates for dominant hand: × = −28; y = −6; z = 58; and non-dominant hand: × = 30; y = −2; z = 48) (Rossiter *et al*., 2013). This was done to facilitate comparisons between different groups. Subsequently, coherence was estimated from the source reconstructed time-series and the EMG data from the active FDI muscle using a similar approach to the one described above. This analysis was performed to quantify both the peak and the area of beta-band coherence in addition to the directionality, i.e. magnitudes of descending (Cortex-to-EMG) and ascending (EMG-to-cortex) components of coherence in the beta band. The latter is dissected by filtering the data with a pre-whitening filter prior to estimating coherence (for a detailed description please see (Halliday, 2015)). This enables a decomposition of descending, ascending and zero-lagged contributions to coherence for each frequency based on the time lag between the two signals. As zero-lagged contribution to coherence may reflect common (non-physiological) activity present in both EMG and EEG data (West *et al*., 2020) this was discarded.

Beta-range coherence area and peak in addition to the magnitude of descending and ascending beta band coherence were extracted. Potential differences due to age group and hand used were then statistically investigated using mixed effect models in Rstudio (v4.0.0). Mixed effects models were chosen as they allow fitting models on incomplete datasets (see Results section). Individual models with the dependent variables were fitted with age group and hand as the independent fixed variable with an interaction (Group × Hand). To account for inter-individual differences in coherence and the repeated measures design we added ‘participants’ as random intercepts. For the analysis of directed coherence (magnitude of descending and ascending coherence components), we added total beta-band coherence area as an additional covariate, as we observed that absolute levels of coherence components scaled with total levels of beta area and beta peak coherence (and co-varied with age group) (Supplementary data S1). Models were fitted using the *lme4* package (Bates *et al*., 2014) and p-values for main and interaction effects were obtained using the *lmerTest* package (Kuznetsova *et al*., 2017) using the Satterthwaite’s method for degrees of freedom. Normality of residuals were checked visually, and log transformations were applied to the dependent variables if this was not fulfilled (this was the case for motor precision; motor variability and the area of beta coherence).

We also quantified the number of individuals displaying significant peaks in corticomuscular coherence in the different age groups and statistically compared these proportions using mixed effect logistic regression fitted using generalized mixed effect models in the *lme4* package with the binomial ‘logit’ family link. Significance of corticomuscular coherence for each individuals was determined based on a NULL hypothesis of uncorrelated signal pairs (Gaussian noise) by computing the upper 95% confidence limit as per (Halliday *et al*., 1995):

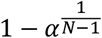

Where α is the confidence limit (0.05) and N is the number of non-overlapping data segments. Presence or absence of significant coherence in the beta-range was binary-coded (1 or 0) from individual coherence spectra for each hand separately. Likelihood-ratio tests were performed for the logistic linear mixed effect models to evaluate significance (Winter, 2013).

For all analyses that involved multiple comparisons we controlled for the false discovery rate (FDR) using the *emmeans* R package. This comprised the group-wise comparisons of (1) motor performance; (2) proportions of individuals displaying coherence; (3) area and peak coherence; (4) descending and ascending magnitude of coherence.

## Results

Participants generally completed the task as intended with both the dominant and non-dominant hand. Some data was excluded due to excessive movements artefacts, an inability to understand and perform the task as intended or technical malfunctions (a total of 17/196 raw data files were excluded; 10 from the dominant hand and 8 from the non-dominant hand). This resulted in a total of 88 EEG-EMG data files that were analyzed from the dominant hand and 90 data files that were used for the analysis for the non-dominant hand. Please also note that two data files from the force-tracing task were not saved following acquisition due to technical issues. Table 1 provides a summary of participant characteristics.

**Table 1:**
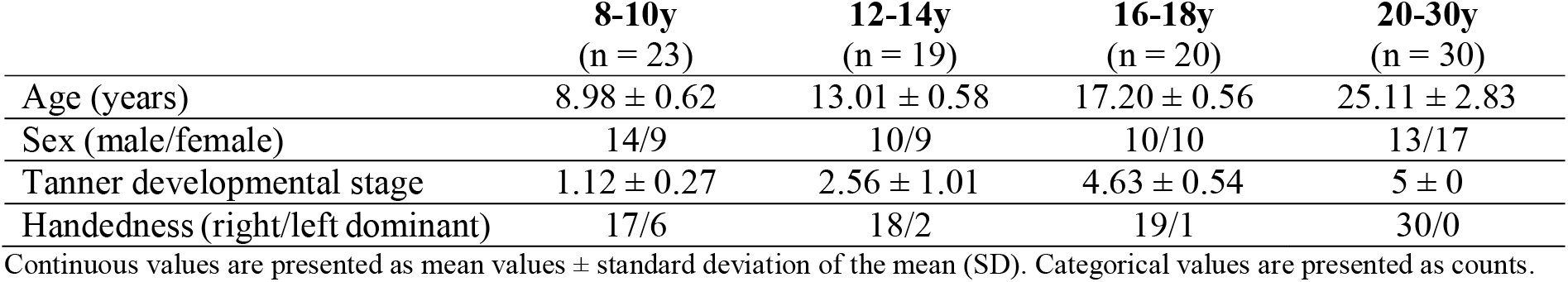
Participant characteristics.

### Differences in performance in precision grip task due to age group and hand used

Participants were instructed to match their force output produced in a precision grip to a horizontal target-line for 2 min. Performance in the tonic precision grip task for the different age groups for both the dominant and non-dominant hand is presented in Figure 2AB. Please note that precision performance for the dominant hand has already been presented elsewhere (Beck et al. in review). For motor precision, a main effect of age group was found (F=21.6; P < 0.001). Subsequent contrasts revealed that individuals in the 20-30y age group significantly outperformed the other groups (β_20-30y vs 8-10y_= 0.44 ± 0.06; P < 0.001; β_20-30y vs 12-14y_=0.25 ±0.06; P < 0.001; β_20-30y vs 16-18y_ = 0.19±0.06; P = 0.007). Additionally, the individuals aged 8-10 also performed significantly worse compared to the 12-14y (β_8-10y vs 12-14y_=-0.19±0.06; P = 0.02) and the 16-18y (β_8-10y vs 16-18y_=-0.25±0.06; P < 0.001). A tendency was observed for the effects of hand used, suggesting that performance was lower for the non-dominant compared to the dominant hand (β_non-dominant vs dominant hand_=-0.03 ± 0.02; P = 0.09). No significant interaction between hand used and age group was found (F=1.20; P =0.31).

**Figure 2:**
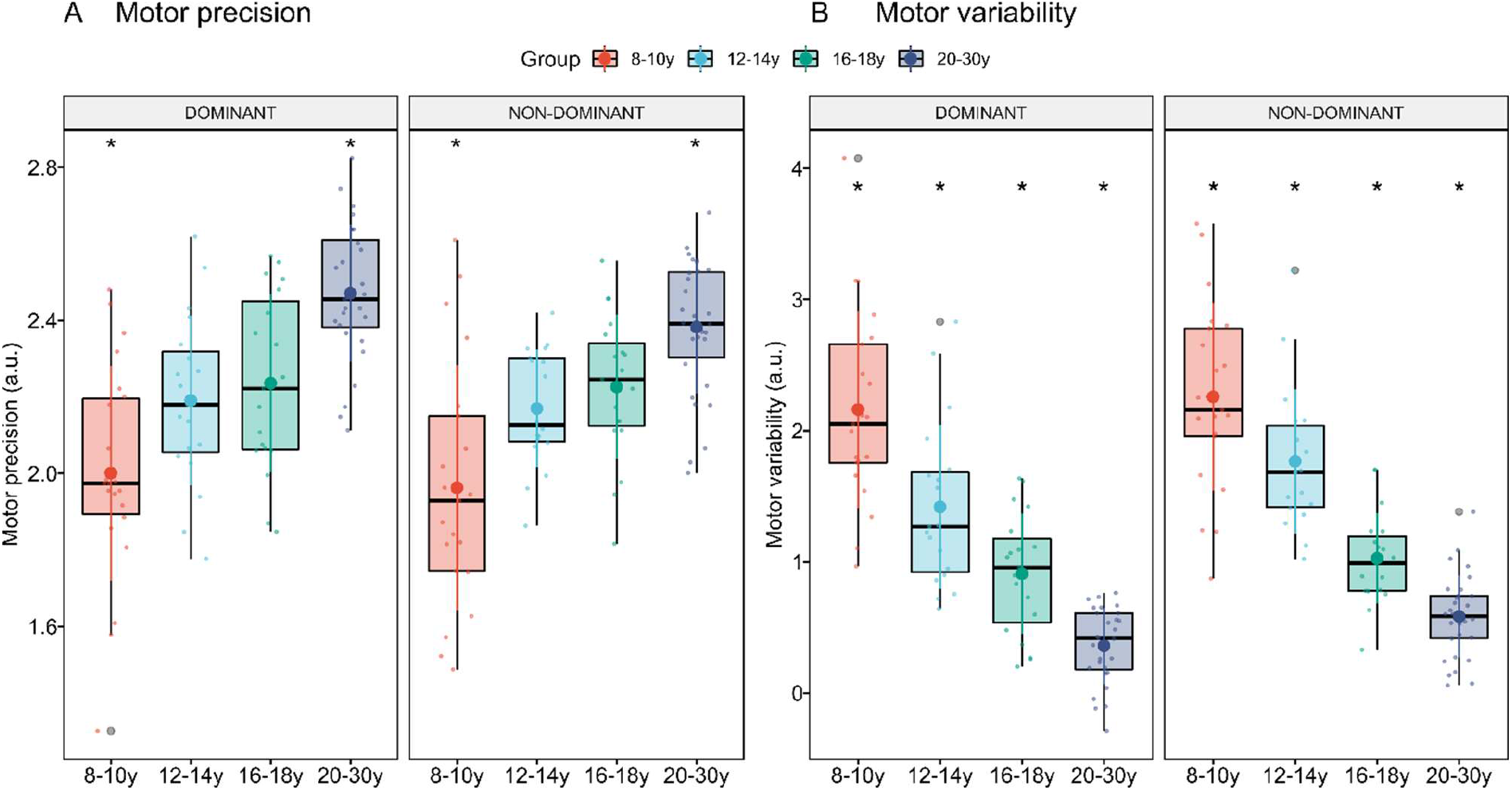
Motor performance. Box plots with individual data points displaying (A) precision performance and (B) motor variability in the force-tracing task for the different age groups divided by hand used (N = 88 for dominant hand; N = 89 for non-dominant hand). Big filled circle reflects mean value and adjoining lines colored by group reflect standard deviations (SD). *signifies a significant difference from the remaining groups averaged over hands (p < 0.05).

For motor variability, a main effect of age group was found (F=64.9; P < 0.001). Subsequent contrasts revealed a linear decrease in motor variability across age groups. Indeed, variability was lower in the 20-30y age group compared to all groups (β_20-30y vs 8-10y_= −1.74 ± 0.12; P < 0.001; β_20-30y vs 12-14y_=-1.15 ±0.14; P < 0.001; β_20-30y vs 16-18y_ = −0.483±0.14; P = < 0.001); lower in the 16-18y age group compared to the 12-14y (β_16-18y vs 12-14y_ = − 0.67 ± 0.15; P < 0.001) and 8-10y (β_16-18y vs 8-10y_ =-1.26 ± 0.15; P < 0.001); And, finally, lower for the 12-14y age group compared to the 8-10y age group β_12-14 vs 8-10y_ =0.59 ± 0.15; P < 0.001). A significant effect was also found for the effects of hand used, driven by a lower degree of motor variability on the dominant hand compared to the non-dominant hand (β_non-dominant vs dominant hand_=-0.18 ± 0.05; P < 0.001). No significant interaction between hand used and age group was found (F=1.11; P = 0.35).

### Differences in corticomuscular coherence due to age group and hand use

To determine the patterns of frequency in corticomuscular coherence across age groups we computed pooled coherence spectra (Figure 1E) and extracted individual peak frequencies of coherence (Figure 1F). In accordance with earlier studies, these descriptive analyses revealed a predominance of beta-range corticomuscular coherence across the entire sample. This led us to restrict our source reconstruction analysis to the beta-band.

To localize brain sources displaying coherent activity in the frequency-domain with the FDI-EMG channel a DICS analysis was performed. The group-level results are shown as volumetric images in Figure 3. Generally, cortical sources displaying maximal coherence with FDI-EMG activity were located in the contralateral hemisphere just anterior or posterior to the central sulcus (left and right precentral or postcentral gyrus; close to M1/S1) for both the dominant and non-dominant hand. Notably, 16-18y and 20-30y were characterized by greater levels of coherence and a slightly more anterior location of cortical sources displaying peak coherence with the FDI activity that extended from precentral into premotor regions (left and right superior frontal gyrus). On the other hand, 8-10y and 12-14y displayed a more posterior locus of cortical peak coherence that extended into the primary somatosensory cortex (S1) and the superior parietal lobule (SPL). It is also important to note that the pattern of coherent cortical sources was more diffuse for children aged 8-10 spreading into the ipsilateral hemisphere, whereas the coherent sources were more spatially focused in older individuals (Figure 3; and supplementary results S2 for 2D density plots of x-y coordinates from sources displaying peak coherence).

**Figure 3:**
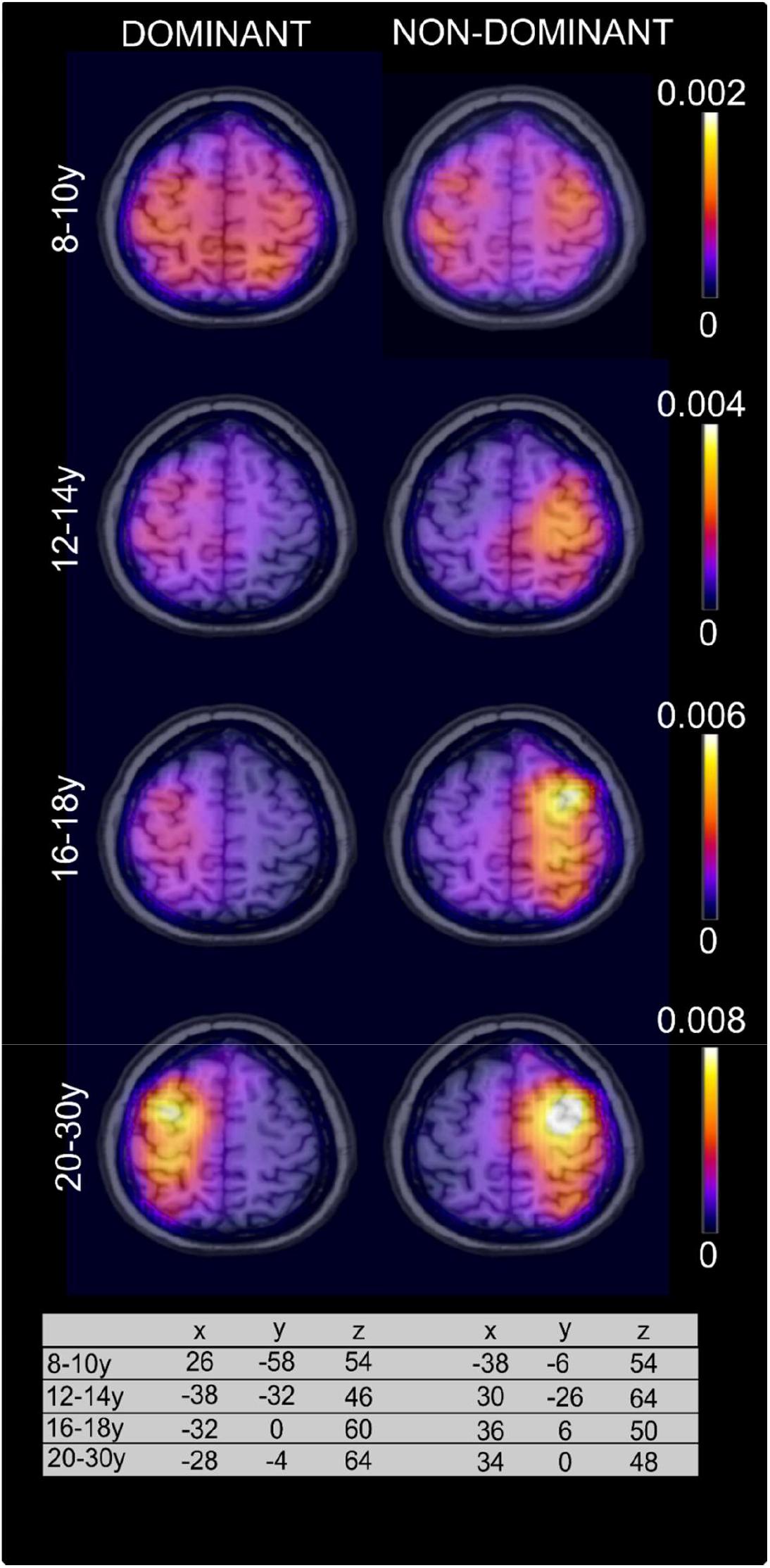
Dynamic imaging of coherent sources. Displays age-related differences in localization of beta-range peak coherence with active FDI muscle for the dominant and non-dominant hand. Sources are displayed on a template MRI provided in the visualization tool MRIcron at z = 58. The coordinates represent the group average location of coherent brain sources in MNI-space (mm). Note that coherence axes are scaled differently for each age group for illustrative purposes.

We quantified the numbers of individuals displaying significant coherence by the upper 95% confidence limit. In Table 2, we present these proportions. Using likelihood-ratio tests, an effect was found for age group (Chi^2^= 19.42; P < 0.001) and hand (Chi^2^ = 16.84; P < 0.001) compared to a null model. The former was driven by larger proportions of 20-30y displaying significant coherence compared to 8-10y (OR_20-30y vs 8-10y_=1.61 ± 0.48; P = 0.0034) and 12-14y (OR_20-30y vs 12-14y_=1.45 ± 0.63; P = 0.008) and a larger proportion of 16-18y displaying significant coherence compared to 8-10y (OR_16-18y vs 8-10y_=1.37± 0.54; P = 0.023) and the 12-14y (OR_16-18y vs 12-14y_=1.21± 0.55; P = 0.04). Furthermore, a larger proportion of individuals displayed coherence on the non-dominant hand compared to the dominant hand (OR=1.43 ± 0.39; P < 0.001).

**Table 2:** Proportion of individuals displaying significant corticomuscular coherence by group.

|  | Dominant hand | Non-dominant hand |
| --- | --- | --- |
| 8-10y | 5/22 | 17/22 |
| 12-14y | 10/19 | 9/18 |
| 16-18y | 10/19 | 20/20 |
| 20-30y | 21/28 | 25/30 |

We compared whether significant differences were present for beta-range coherence area as a function of age group and hand. This revealed an effect of age group (F = 4.20; P = 0.008). As can be seen in figure 4A, the effect was driven by greater levels of beta-range coherence in the 20-30y group compared to the 8-10y group (β_20-30y vs 8-10y_=0.30 ± 0.09; P = 0.008) and by a tendency towards greater levels of coherence in the 20-30y group compared to the 12-14y group (β_20-30y vs 14-12y_ =0.23 ± 0.10; P = 0.059). Furthermore, a strong tendency towards a main effect of hand was also found (F = 3.87; P = 0.052), suggesting that coherence magnitudes were larger for the non-dominant compared to the dominant hand across groups (β_ND vs DOM_=0.106 ± 0.05). No interaction between group and hand-used was found (F = 1.18; P = 0.32).

**Figure 4:**
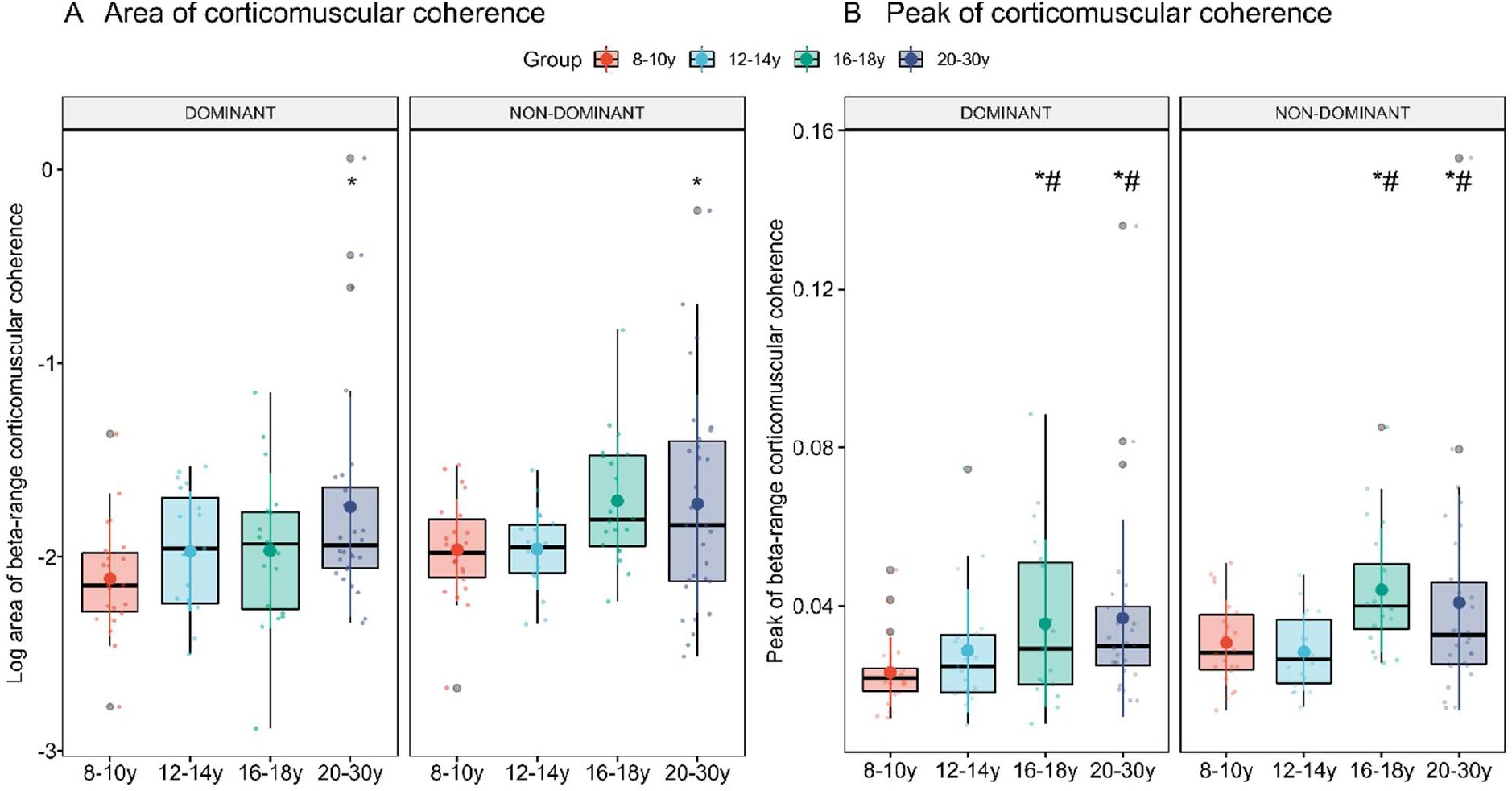
Beta-band corticomuscular coherence. Box plots with individual data points and means displaying age-related differences in the log area (A) and peak (B) of beta-band corticomusclar coherence between cortical source and active FDI muscles on the dominant and non-dominant hand. Statistically significant differences from the 8-10y group and the 12-14y group are marked by * and #, respectively (p < 0.05).

Comparable results were found for beta-range peak coherence: A main effect of age group was observed (F = 4.86; P = 0.004) that was driven by greater peak coherence in the 20-30y group compared to the 8-10y (β_20-30y vs 8-10y_ =0.012 ± 0.004; P = 0.015) and the 12-14y (β_20-30y vs 12-14y_ =0.010 ± 0.004; P = 0.027), and by greater peak coherence in the 16-18y group compared to the 8-10y (β_16-18y vs 8-10y_ = 0.013 ± 0.004; P = 0.015) and the 12-14y (β_16-18y vs 12-14y_ = 0.011 ± 0.004; P = 0.026)(Figure 4B). A tendency was found for the main effect of hand (F = 3.26; P = 0.074), that was driven by marginally greater levels of peak coherence on the non-dominant compared to the dominant hand (β_ND vs DOM_=0.005 ± 0.003). No interaction between group and hand was found (F = 0.50; P = 0.68). Notably, similar age-related effects were seen when comparing peak coherence at the individual peak location of coherence (See supplementary results, Figure S3), suggesting that it is unlikely that developmental differences were systematically influenced by the fact that sources were extracted from the grand-averaged peak location.

For directionality components, we found a significant effect of age group on the levels of descending (Cortex-to-EMG) coherence (F = 3.56; P = 0.017). Specifically, individuals in the 20-30y group had significantly larger magnitudes of descending coherence compared to the 8-10y group (β_20-30y vs 8-10y_=0.003 ± 0.001; P = 0.031) and the 12-14y group (β_20-30y vs 14-12y_=0.003 ± 0.001; P = 0.039), while a tendency towards greater magnitudes was found between the 20-30y and the 16-18y was found (β_20-30y vs 16-18y_=0.003 ± 0.001; P = 0.10)(Figure 5A). Magnitudes of descending and ascending coherence were found to scale with total levels of coherence (see supplementary results S1). Therefore, we added total levels of coherence (indexed as either log area and peak coherence) as covariates in the mixed model. The main effect of age group persisted when controlling for peak beta-range coherence (F = 4.53 = 0.005), but it was reduced to a tendency when controlling for beta-range coherence area (F = 2.00, P = 0.12). No main effect of hand was found (F 2.47; P = 0.12), but levels were greater for the non-dominant hand compared to the dominant hand. No interaction between age group and hand was observed (F = 0.09; P = 0.96). For the ascending component, we found an effect of age group (F=2.75; P = 0.048), but this was no longer seen when controlling for the total area (P = 0.98) or peak (P = 0.73) beta-range coherence (therefore we refrain from performing pairwise comparisons). No effects of hand (F = 0.005; P = 0.95) or interactions between age group and hand (F = 1.75; P = 0.16) were observed (Figure 5B).

**Fig 5.**
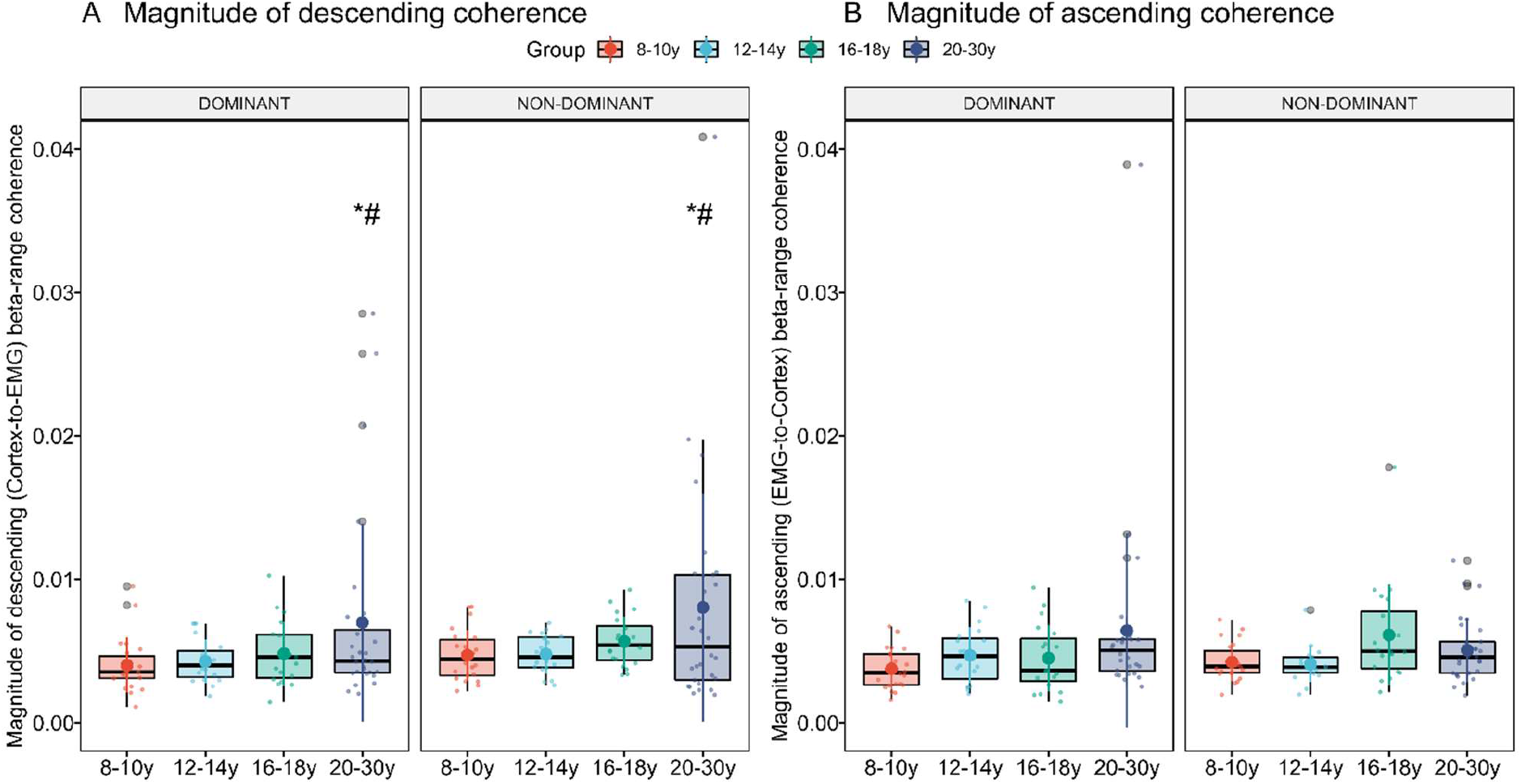
Directed corticomuscular coherence. Box plots with individual data points and mean level displaying age-related differences in the descending (A) and ascending (B) amplitudes of beta-band corticomusclar coherence between cortical source and active FDI muscles on the dominant and non-dominant hand. Statistical significant differences from the 8-10y group and the 12-14y group is marked by * and #, respectively (p < 0.05).

As a final step, we used regression models to determine potential statistical dependencies between measures of beta-range coherence and motor performance (quantified as motor precision and motor variability). We did not find any significant associations between the included measures of coherence and motor precision (beta-range area: β_Beta-range area_=0.014 ± 0.036, P = 0.70; beta-range peak: β_Beta-range peak_= −0.13 ± 0.75, P = 0.86; magnitude of descending coherence: β_Beta-range descending_=-0.19 ± 3.84, P = 1; magnitude of ascending coherence: β_Beta-range ascending_= 6.50 ± 3.95, P = 0.10). Additionally, no significant associations were found for motor variability (beta-range area: β_Beta-range area_=-1.54 ± 1.01, P = 0.13; beta-range peak: β_Beta-range peak_= −35.6 ± 20.8, P = 0.090; magnitude of descending coherence: β_Beta-range descending_=-113.0 ± 104.4, P = 0.28; magnitude of ascending coherence: β_Beta-range ascending_= 102.3 ± 112.5, P = 0.37).

## Discussion

We studied developmental differences in corticomuscular coherence during a visually guided precision grip force-tracing task using source-reconstructed brain activity from EEG data. The proportion of individuals displaying significant beta-range corticomuscular coherence and magnitude of beta-band corticomuscular coherence was found to be greater for older adolescents and adults compared to younger adolescents and children. This was paralleled by greater magnitudes of descending – but not ascending - connectivity for adults compared to children. We further observed that adults and older adolescents exhibited a more anterior cortical locus of peak coherence compared to children and young adolescents. Lastly, we observed that corticomuscular coherence was greater for the non-dominant compared to the dominant hand. Our study provides a detailed characterization of developmental similarities and differences in corticomuscular connectivity during control of low-level force output from the intrinsic hand muscles in typically developed children, adolescents and adults. By demonstrating differences in corticomuscular connectivity between the non-dominant and dominant hand, our results highlight yet another experimental feature that affects corticomuscular coupling. This has potential implications for the comparison of results between previous studies as well as for the understanding of the behavioral relevance of beta-band synchrony in the corticomuscular system.

Comparing corticomuscular coherence in children, adolescents and adults revealed several interesting findings. Characterizing the distribution of peak coherence at the sensor-level for different age groups revealed a similar functional organization of oscillatory coupling within the corticomuscular system. That is, during the tonic precision grip task, children, adolescents and adults predominantly displayed peak EEG-EMG coherence in the beta range (15-30Hz). This finding, and earlier evidence of increased beta-range coupling during tonic contractions in humans and monkeys (Baker, 2007), led us to restrict the DICS source localization procedures to the beta range. Subsequent detected levels of peak cortical source-EMG coherence were generally quite low (range 0.001 – 0.15), but still within the range commonly reported for corticomuscular coherence in both humans (on the sensor-level; e.g James et al., 2008 for forearm extensors) and monkeys (using LFPs; e.g. Baker et al., 1997 or Baker et al., 2003).

The proportion of individuals displaying significant coherence on either their dominant or non-dominant hand amounted to 66% for the entire sample. This percentage is slightly higher than reported previously (e.g. James et al., 2008; Ushiyama et al., 2011), but this probably reflects differences in the approach taken to evaluate significance of coherence as well as differences the experimental design; e.g. whether the dominant or non-dominant limb was used (see below) or whether the analyses were based on source or sensor data. More interestingly in the context of developmental differences, we found that older adolescents (16-18y) and adults (20-30y) displayed significant corticomuscular coherence more frequently than children (8-10y) and young adolescents (12-14y). In children younger than 10y, the proportion of individuals displaying beta-range coherence was generally low (especially for the dominant hand), which could indicate that the drive from corticospinal cells to motoneurones likely not display prominent oscillatory components in late childhood (∼10y). These results are similar to the study by James and co-workers (James *et al*., 2008). As was the case with the results presented here, they found little evidence of clear corticomuscular coherence in children before the age of 12y.

In a similar vein, we found that adults and older adolescents were characterized by displaying greater amplitudes of corticomuscular coherence. This was expressed both as differences in the area of beta-range coherence and in the peak values of coherence. These results from a large sample of typically developing individuals are well in line with previous studies demonstrating age-related differences in levels of functional cortico-motor coupling in children, adolescents and adults (James *et al*., 2008; Graziadio *et al*., 2010; Spedden *et al*., 2019*b*), likely reflecting maturational differences in the oscillatory interactions between cortical and muscle activity during tasks requiring accurate control of a tonic contraction. Taken together, these results suggest that oscillatory corticomuscular coupling emerges after 10y of age and progressively strengthens during adolescence until early adulthood. These developmental profiles of functional connectivity parallel the ones reported for structural changes within and between cortical structures and in ascending and descending white matter tracts (Paus *et al*., 1999; Lebel & Beaulieu, 2011), suggesting that the two interact (Uhlhaas *et al*., 2010). Earlier studies have demonstrated that coherence is transiently (and oppositely) modulated by skilled motor practice (Perez *et al*., 2006; Larsen *et al*., 2016) and limb immobilization (Lundbye-Jensen and Nielsen, 2008), suggesting that the corticomuscular system adapts in the face of changes in sensorimotor demands, i.e. with learning or disuse, which can manifest as brief changes in oscillatory interactions. Here, we demonstrate developmental differences in functional coupling that could reflect adaptation and tuning of sensorimotor processes occurring over longer time scales. The increased levels of corticomuscular synchrony could be interpreted as reflecting a greater or more efficient engagement of oscillatory control of corticospinal networks during the control of low-level force output in old adolescents and adults compared to children (Baker *et al*., 1999; Schoffelen *et al*., 2005).

We also explored potential age-related differences in the location of brain sources displaying peak coherence with the active FDI muscle using beamforming techniques. Peak coherence was generally observed for sources located around the central sulcus in the contralateral cortical hemisphere. This has been robustly reported previously for tasks using the hand and forearm muscles in adult humans (Gross *et al*., 2000; Kilner *et al*., 2000). Specifically, this spatial organization was observed in individuals aged 12 and above, whereas more spatially diffuse patterns of coherence were seen for children younger than 10 years old extending into the ipsilateral hemisphere. The diffuse spatial patterns of coherence might reflect that very low levels of coherence were generally observed in the young children. Another interesting observation was that average peak coherence was more focused and seen at a slightly more anterior site from late-adolescence and onwards compared to their younger counterparts. Specifically, peak coherence was found in the precentral gyrus extending into premotor regions from 16y and onwards, whereas younger individuals (8-10y and 12-14y) were characterized by displaying peak coherence in regions close to the postcentral gyrus (around primary somatosensory cortex (S1) extending into parietal regions). Corticomuscular coherence has traditionally been considered a motor phenomenon reflecting oscillatory activity descending from the M1 via corticospinal pathways (Hansen & Nielsen, 2004; Gerloff *et al*., 2006), but growing evidence suggests that ascending sensory activity also contributes to the coupling between sensorimotor brain regions and muscle (Riddle & Baker, 2005; Witham *et al*., 2011). Coherent activity between cortex and muscle might thus rather reflect sensorimotor integration processes, constituting a loop in which ascending sensory information (feedback) interact with descending motor activity (feedforward). Previous studies have suggested that spatiotemporal fluctuations in coherence during steady-state motor tasks reflect the underlying dynamic interactions within this sensorimotor loop (Graziadio *et al*., 2010; Maezawa *et al*., 2016). Specifically, Graziadio et al., (2010) argued that coherence between fronto-central electrodes and EMG mainly reflected feedforward (motor) processing, whereas parieto-central coherence with EMG chiefly revealed feedback (sensory) processing. It could be speculated that the age-related differences in spatial organization of coherence in the present study could reflect differences in the relative reliance on these processes. That is, the more anterior cortical site of peak coherence observed in older adolescents and adults might suggest a predominance of feedforward (descending) control, whereas the more posterior location of peak coherence in younger individuals would suggest a greater role of feedback (ascending) control.

We also used measures of directed corticomuscular coherence to formally tease apart the contribution of descending and ascending flow of information. Absolute levels of descending (cortex-to-muscle) coherence was greater in adults compared to children, and these age-related differences persisted after controlling for total amount of beta-range coherence (indexed as peak values of corticomuscular coherence). This was not the case for the ascending components of coherence. But how might we understand these age-related differences?

There is convincing evidence showing that our ability to perform movements accurately and smoothly relies on constructed internal models of the body and how it interacts with the environment. A central idea is that the CNS uses these models to predict the sensory consequences of intended actions. In doing so, internal models aid in ensuring that movements are performed as intended, and are involved in correcting ongoing and optimizing future movements when this is not the case (Flanagan *et al*., 2006; Wolpert & Flanagan, 2016). These processes kick in when a mismatch arises between the predicted and the actual sensory consequences of action. This triggers rapid corrective responses that attempts to adjust the movement to fit the intentions (Johansson & Westling, 1988), and acts as a learning signal (i.e. a sensory prediction error) that can be used to optimize central nervous motor programs of subsequent movements (Shadmehr *et al*., 2010; Wolpert & Flanagan, 2016). These (motor) learning processes may occur on a daily basis throughout the lifespan, e.g. as we dexterously manipulate objects. This may lead to more elaborate and robust internal models, and hence more exact predictions of the sensory consequences of ones actions. With time, this could entail a greater use of (precise) feedforward (predictive) control and a concomitant gradual reduction in the importance and necessity of adjustments based on sensory cues (i.e. feedback) (Moran *et al*., 2014) during motor actions, leading to a shift in the relative reliance on feedforward and feedback processing as children become adults. Our results could be interpreted in this sense: A greater degree of descending connectivity was reported for adults compared to children and taken together with the posterior-to-anterior shift in the cortical site of peak coherence observed from 16y and onwards this could represent an increased reliance on feedforward and predictive control during development. Earlier studies have demonstrated similar developmental shifts using measures of directed (or effective) connectivity at the cortical level during tasks requiring response inhibition (Hwang *et al*., 2010) and force control (Beck et al., in review), but also for directed corticomuscular connectivity during a force-tracing task involving the ankle muscles (Spedden *et al*., 2019*b*). Additionally, during movement tasks such as walking, but also while performing precision grips, an increase in feedforward control has been reported from early to late childhood (Forssberg *et al*., 1992; Lorentzen *et al*., 2019), while decreases in task-related transmission from sensory afferents to motor neurons has been demonstrated to occur at the spinal level concomitantly (Hodapp *et al*., 2007).

The implications of these developmental differences in neural control strategies remain unclear. The functional relevance of corticomuscular connectivity to behavior has been extensively debated in the past decades. Beta band coherence is prominent during tasks requiring maintained motor outputs (Baker *et al*., 1997), but decreases in magnitude or disappears with actual movement (Baker *et al*., 1997; Omlor *et al*., 2007; Mehrkanoon *et al*., 2014). This has led to the suggestion that oscillatory activity in the beta-range is involved in maintaining steady motor output (Engel & Fries, 2010; van Wijk *et al*., 2012). We found greater levels of beta-band coherence in adults who were also characterized by performing the task with greater precision and less variability. It therefore seems reasonable to speculate that levels of corticomuscular coherence are associated with the ability to precisely control static force levels. This is in fact also supported by earlier studies showing greater magnitudes of beta-band coherence when precision (or requirements for precision) in a motor task is high (or variability of motor output is low) (Kristeva *et al*., 2007; Witte *et al*., 2007; Mendez-Balbuena *et al*., 2012). In contrast, no clear statistical associations were found between measures of coherence and precision grip performance in the present study. This could be related to the substantial variability inherent in measures of coherence between individuals (as seen in the present study, but also specifically investigated in Ushiyama et al., 2011), all of whom successfully performed the task. Perhaps our task was not sufficiently demanding to detect associations between motor performance and beta coherence.

Another interesting observation was that levels of corticomuscular coherence generally seemed larger and more frequent on the non-dominant hand compared to the dominant hand. Ushiyama and co-workers (2010) previously reported that corticomuscular coherence was greater for lower-limb muscles compared to upper-limb muscles; and for distal muscles compared to the proximal muscles. Here, we report results suggestive of small between-limb differences in corticomuscular coherence for the same muscle due to limb dominance, with larger levels of coherence reported for the non-dominant compared to the dominant hand. Notably, these effects were most likely not due to order effects alone (i.e. that participants in the main experiment always performed the task with the non-dominant hand first) as suggested by a set of control experiments in which the order was reversed, but the pattern remained (see supplementary results S4). This suggests a task-related modulation of coherence amplitudes by alternating the hand used. We speculate that the difference in coherence due to handedness could be related to the additional demands for controlling pinch force imposed by using the non-dominant hand. The fact that precision performance was lower and variability was larger on the non-dominant hand compared to the dominant could indeed indicate that the task was more challenging when it was performed with the non-dominant hand. This might have required the use of a more efficient (or even an alternative) neural strategy for controlling and adjusting force production during this state. The corticomuscular system might attempt to overcome this challenge by tuning the coupling efficiency between the cortex and the muscles, which may allow a more efficient binding of sensory and motor information. This would be evident as an increased magnitude of corticomuscular coherence. In line with this idea, greater coherence amplitudes have previously been observed during dual-task conditions (Perez *et al*., 2012) and in visually-guided walking demanding accurate placement of the feet based on visual cues as opposed to normal walking without such requirements (Spedden *et al*., 2019*a*). It could be speculated that the increased levels of corticomuscular coherence could also be interpreted as reflecting a greater corticospinal involvement when the nervous system is faced with more task-related complexity. For generalization purposes, it would be of further interest to investigate whether the reported differences due to limb dominance can be extrapolated to the cortical control of other muscles. In any way, these differences warrant careful attention to the experimental procedures when comparing results between studies.

Ontogenetic development of fine motor skills follows a protracted trajectory, but the underlying neural mechanisms are not well characterized. Here, we demonstrate that oscillatory corticomuscular connectivity emerges after 10y of age and is more prevalent and greater in magnitude in adults and older adolescents (from 16y and onwards) compared to younger adolescents and children (14y and younger). We further found that adults were characterized by greater levels of descending connectivity in the corticomuscular system. This was paralleled by the finding of a posterior-to-anterior shift in the location of cortical sources displaying peak corticomuscular coherence in the contralateral hemisphere from late adolescence. Collectively, these results suggest that task-related functional corticomuscular connectivity continues to develop through adolescence. We further found small differences in corticomuscular coupling between the dominant and non-dominant hand, with larger amplitudes of coherence observed for the non-dominant hand, suggestive of a task-related modulation of corticomuscular coupling within individuals that may be related to task difficulty. It also revealed yet another factor that researchers have to consider when designing new studies and comparing results between studies that are already available. Collectively, our results help us better understand developmental differences in the corticomuscular control of fine movements in children, adolescents and adults.

## Acknowledgements

We would like to thank the participants and their parents for their participation. Furthermore, we would like to thank Gitte Abrahamsen and Frederikke Toft Kristensen for assisting in the data acquisition.

## Supplementary results

**S1 Associations between log beta area corticomuscular coherence and amount of descending (cortex-to-EMG) and ascending (EMG-to-cortex) components**.

Figure S1 below presents the associations between the area of corticomuscular coherence in the beta-range and the decomposed amount of descending (Cortex-EMG; red) and ascending (EMG-Cortex; blue) beta-range coherence. Pearson correlation coefficients and corresponding p-values for the correlation analyses are displayed for both the dominant and non-dominant hand.

**Figure S1.**
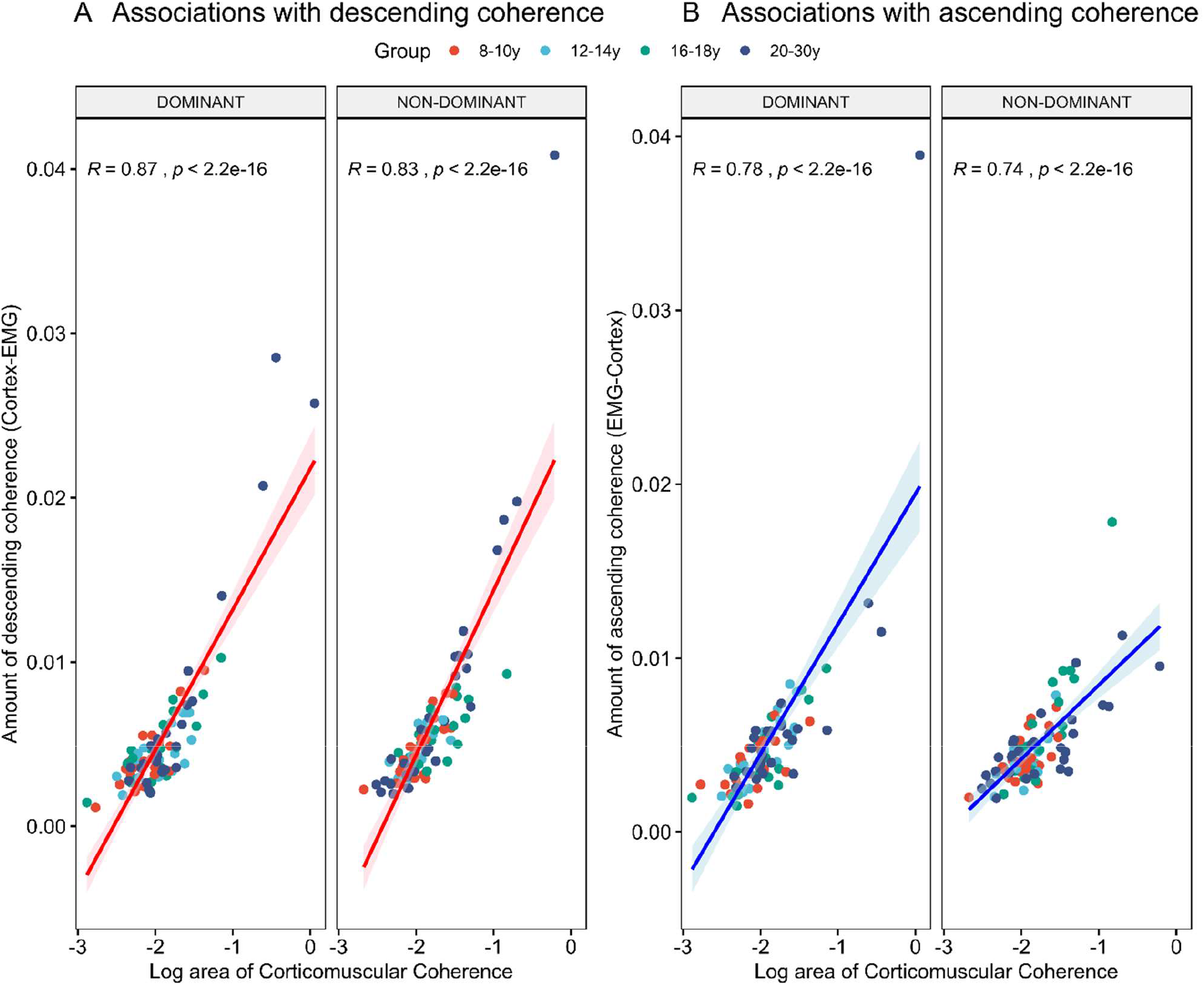
Associations between log area of corticomuscular coherence and amount of descending (left) and ascending (right) coherence for both the dominant and non-dominant hand.

**S2 Distribution of brain sources displaying coherence with muscle activity**.

Below we provide 2D density plots of peak locations of sources displaying coherence with muscle activity for both the dominant and non-dominant hand.

**Fig S2.**
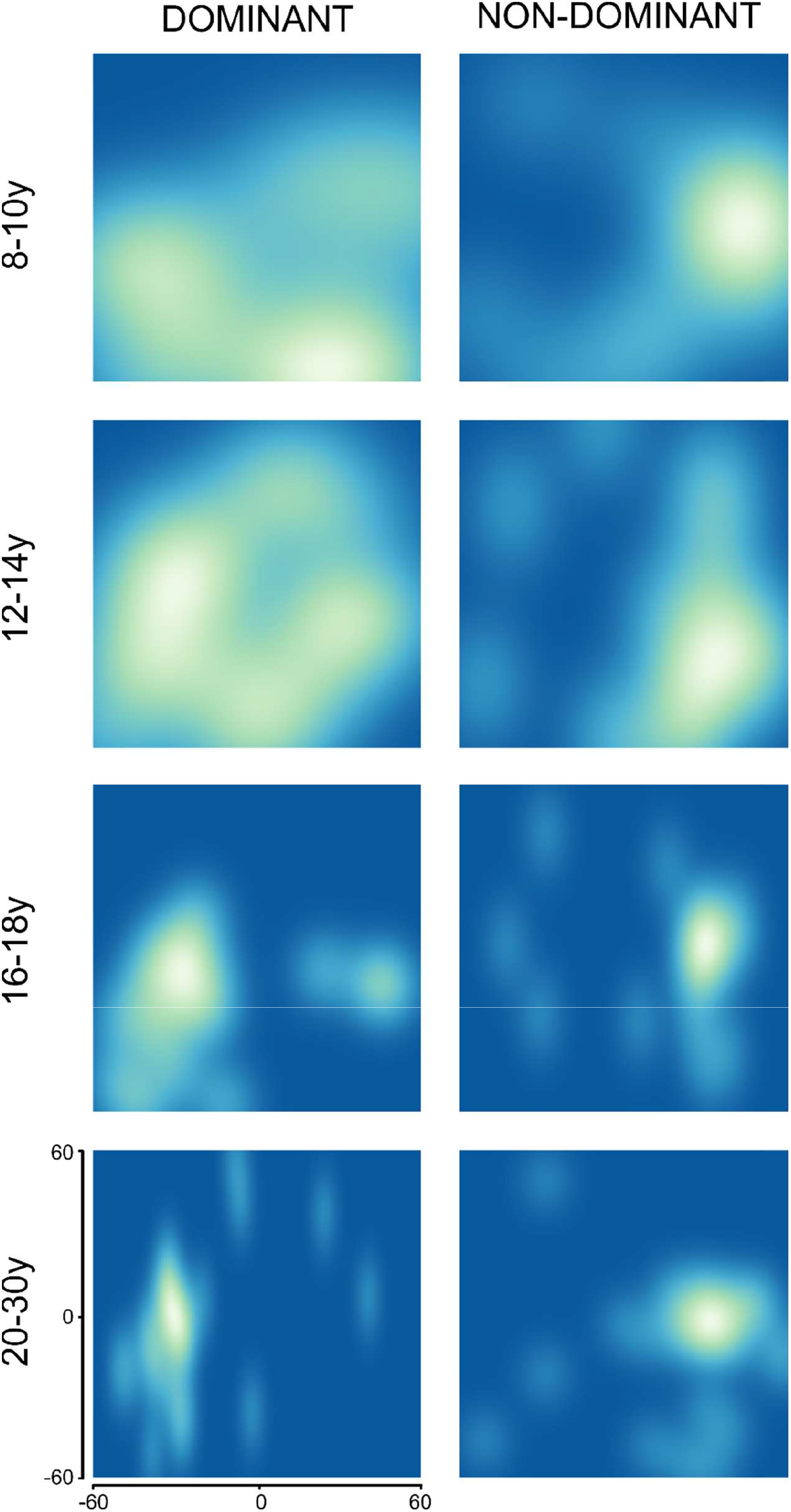
2D density plot of sources displaying peak coherence with active FDI muscle in MNI space. X-axis span left-to-right with x=0 representing the midline. Negative values are in the left hemisphere, whereas positive values reflect the right hemisphere. Y-axis reflect the posterior-anterior direction. Note the diffuse localization of cortical sources in the youngest individuals (especially for the dominant hand).

**S3 Developmental differences in corticomuscular coherence at individual peak coherence**

In the main analysis, we used the sample average location of peak corticomuscular coherence to study developmental differences in corticomuscular coherence. These differences in corticomuscular coherence could potentially be confounded by differences in the spatial distribution of sources displaying coherent activity with the active muscle. Therefore, we tested whether age-related differences in magnitudes of coherence were present using the individual peak values from the DICS analysis. This analysis revealed a significant effect of group (F = 3.94, P = 0.011), and subsequent pairwise comparisons suggested that this was driven by 20-30y displaying significantly more corticomuscular coherence than 8-10y (β_20-30y vs 8-10y_ = 0.015 ± 0.005; p = 0.012) and the 12-14y (β_20-30y vs 12-14y_=0.012 ± 0.005; p = 0.054) (Figure S). These results parallel the ones reported in the main analysis.

**Figure S3.**
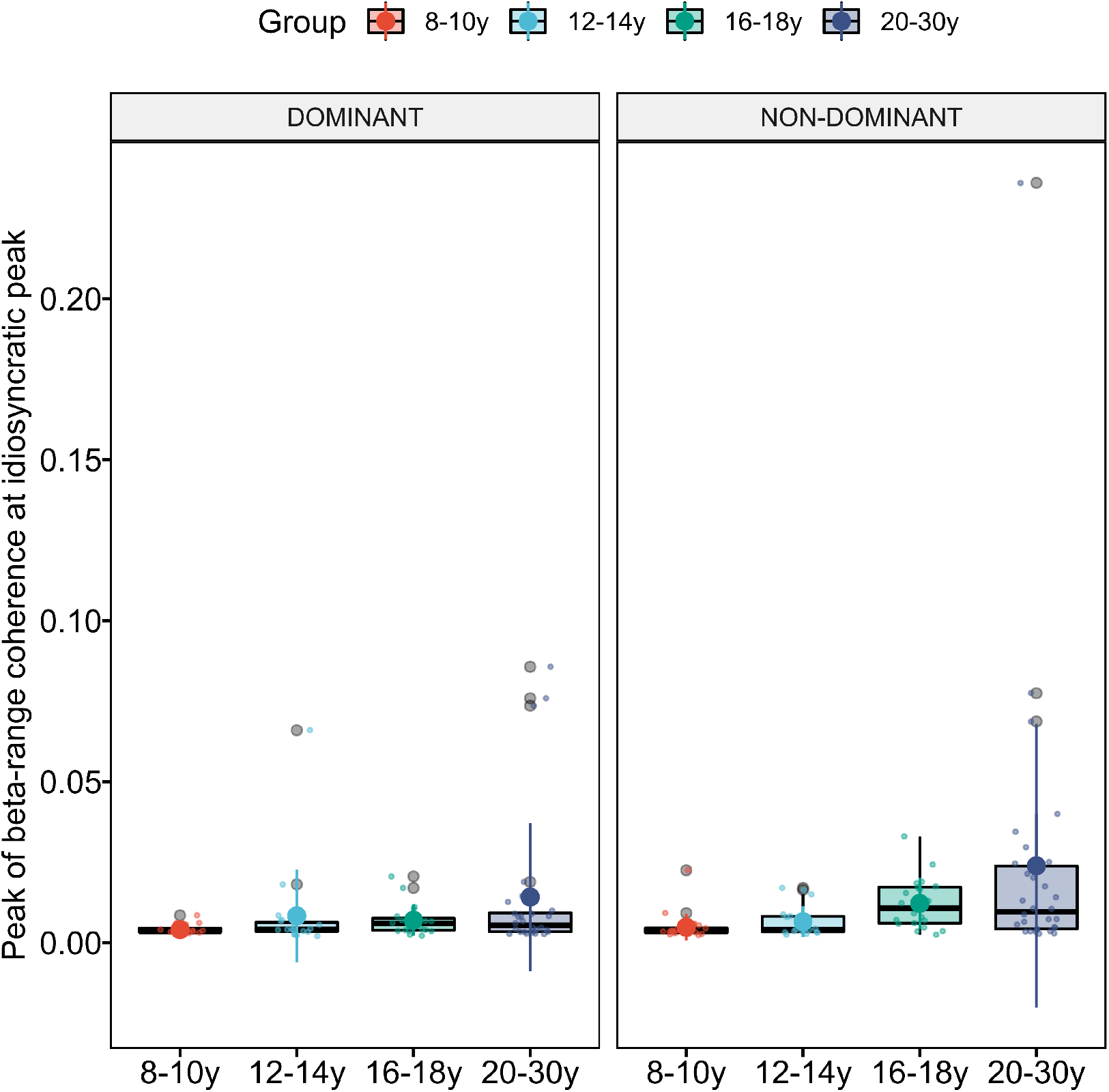
Peak of corticomuscular coherence at individual peak locations from dynamic imaging of coherent sources (DICS).

**S4 Differences in corticomuscular coherence in the dominant and non-dominant hand: effects of order of testing**

We also tested whether the trends towards differences due to hand dominance were contingent upon order of testing. That is, individuals in the main experiment always performed the task with the non-dominant hand first, and it could be speculated that differences in coherence could relate to novelty of the task. To control for potential order-effects on comparisons between the dominant and non-dominant hand, we obtained data from an additional 15 adult individuals (23.5 ± 1.6 years old; 8 females) that did not participate in the main experiment and were naïve to the task presented. For these individuals, the order of tests was flipped so that they performed the task with the dominant hand first. Adult individuals were chosen for the control experiment as they (most) frequently displayed significant corticomuscular coherence. The analysis was performed on sensor level data and time-series were extracted from the C3 and C4 electrodes to compute corticomuscular coherence with the FDI muscle activity on the dominant and non-dominant hand, respectively.

**Fig S4.**
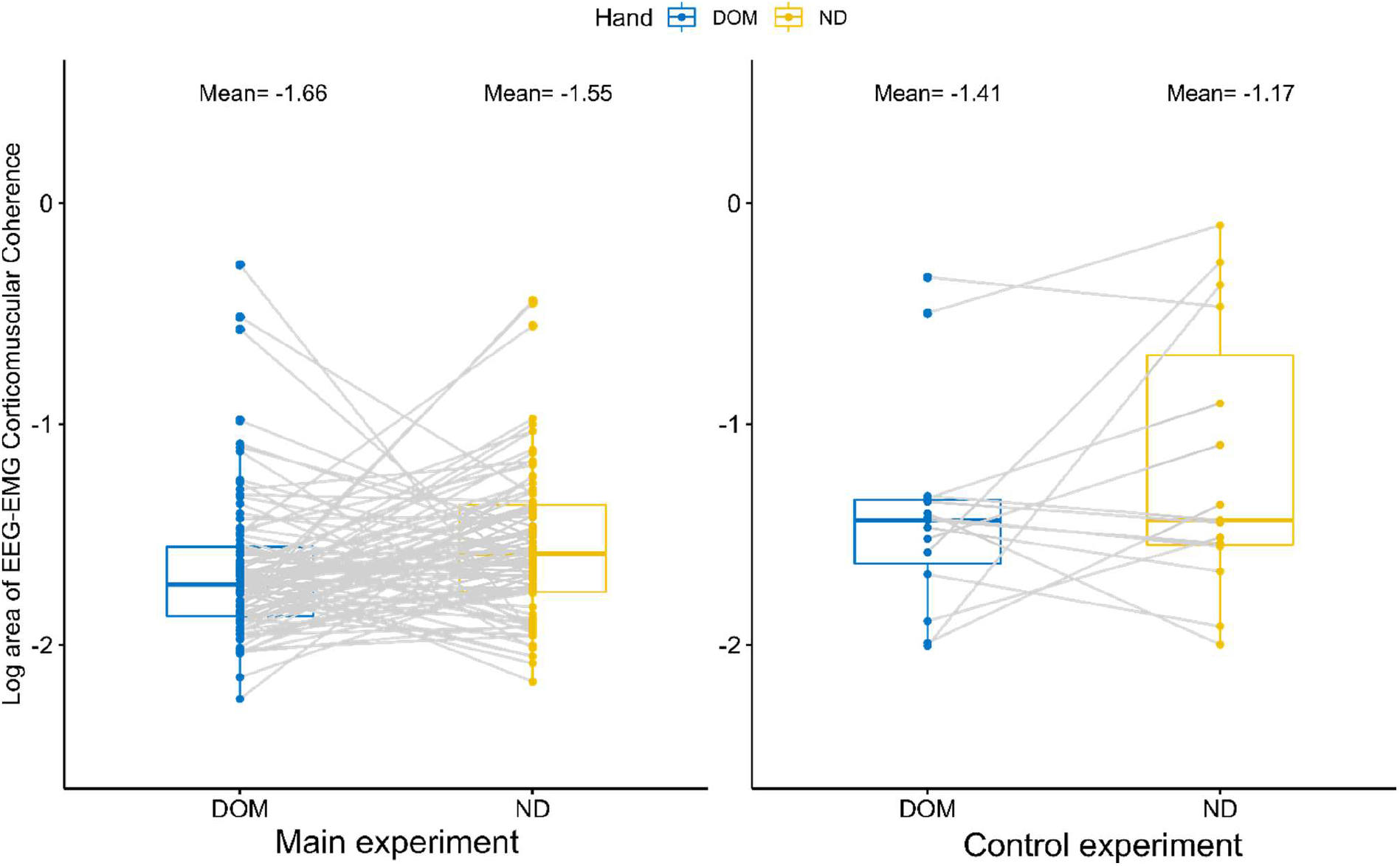
Corticomuscular coherence for the dominant and non-dominant hand with the reverse order of testing (dominant before non-dominant hand). DOM = dominant hand; ND = non-dominant hand.

Differences in corticomuscular coherence due to hand use seemed to exhibit a similar pattern in the control sessions as in the main experiment. As such, a tendency towards greater EEG-EMG corticomuscular coherence was found from a paired t-test for the non-dominant compared to the dominant hand in the control experiment (t(14)=1.44; P = 0.17). This suggests that differences due to hand dominance are likely not exclusively related to the order by which the tests were performed.

